# Perceptions, attitudes and practices towards scabies in communities on the Bijagós Islands, Guinea-Bissau

**DOI:** 10.1101/574327

**Authors:** Maria João Lopes, Eunice Teixeira da Silva, Janete Ca, Adriana Gonçalves, Amabelia Rodrigues, Cristóvão Manjuba, Jose Nakutum, Umberto D’Alessandro, Jane Achan, James Logan, Robin Bailey, Anna Last, Steve Walker, Michael Marks

**Affiliations:** Clinical Research Department, Faculty of Infectious and Tropical Diseases, London School of Hygiene & Tropical Medicine, London, United Kingdom; Infectious Diseases Department, Hospital Prof. Doutor Fernando Fonseca, Amadora, Portugal; Region Sanitaria Bolama-Bijagós, Bubaque, Guinea Bissau; Bandim Health Project, Guinea Bissau; Ministry of Public Health, Guinea Bissau; MRC The Gambia at the London School of Hygiene & Tropical Medicine; Disease Control Department, Faculty of Infectious and Tropical Diseases, London School of Hygiene & Tropical Medicine, London, United Kingdom; Hospital for Tropical Diseases, London, United Kingdom

## Abstract

**Introduction:** Scabies is highly endemic among impoverished populations and has been recently included in the WHO’s list of neglected tropical diseases (NTDs). Community support and behavioural changes are essential for the success of control interventions. This study aimed to explore beliefs, prevention attitudes and health care-seeking behaviours towards scabies in the Bijagós Archipelago of Guinea-Bissau.

**Methods:** Data were collected through two methods. Community key informants (community members, community health workers, healthcare workers and traditional healers) were interviewed using snowball sampling. A questionnaire covering perceptions, attitudes and practices was administered to community members using random cluster sampling. Thematic analysis of qualitative data was applied to identify themes. Descriptive statistics were used for quantitative data analysis.

**Results:** There was a satisfactory awareness about scabies, but perceptions about disease causation and transmission were imprecise. Misconceptions about personal hygiene as the primary measure for scabies prevention were recurrent. Some participants recognised the importance of early treatment to interrupt transmission. Treatment of close contacts was not considered important. Costs were the main determining factor for treatment choice between traditional healer and the local health centre. Late presentation and delayed treatment were common and associated with poverty and stigmatisation. Scabies impaired quality of life by affecting social interactions, health, fitness to work and school attendance.

**Conclusions:** There is a need to improve education, recognition, management and affordable access to treatment. Community education, healthcare workers’ training and skin NTDs integrated control programmes should address the challenges highlighted in this study.

**Authors Summary:** Scabies is a common skin infection in low income settings. We conducted a study in Guinea-Bissau to explore the knowledge, attitudes and practices about scabies. We conducted interviews with healthcare workers, traditional healers and community members and additionally used an oral-administered questionnaire with a larger sample of community residents. Most individuals had knowledge of scabies and were aware that person to person transmission occurred. However personal and environmental hygiene were both incorrectly identified as particularly important in the transmission of scabies. Cost played a major role in determining where individuals sought care and both poverty and disease associated stigma resulted in delays seeking care. There is a need to improve community and health care worker education about scabies and improve affordable access to treatment.

## Introduction

Scabies, caused by infestation with the mite, *Sarcoptes scabiei* var. *hominis*, is a parasitic skin infestation, with a worldwide distribution, which is believed to affect up to 200 million people globally, particularly impoverished populations in low-income settings.^1,2^ In tropical settings, the average prevalence in children is between 5 and 10%^3^ and unpublished data in the Bijagós Islands communities estimated a scabies prevalence of around 5% during the dry season (M Marks, personal communication)..

The primary clinical manifestations are skin lesions (erythematous papules, pustules, small burrows and tunnels) and intense pruritus.^4^ Scabies complications are commonly underestimated. The direct consequences of scabies include an impact on quality of life, due to sleep disturbances caused by pruritus, and stigma, discrimination and psychological distress.^3^ Infestation may influence health-seeking behaviours and treatment compliance.^4,5^ In resource-poor countries scabies is also associated with an increased burden of secondary skin infections.^3^ Infections with Group A Streptococci may lead to acute post-streptococcal glomerulonephritis and acute rheumatic fever.^6,7^

Continued scabies transmission within communities is related to delayed treatment, as a result of delayed diagnosis, poor access to treatment, non-adherence with treatment or a failure to treat household contacts.^4,8^

In 2017, the World Health Organization classified scabies as a Neglected Tropical Disease (NTD) in recognition of the burden and impact of the disease globally. There is increasing interest in mass drug administration (MDA) with ivermectin as a control strategy for scabies in highly endemic settings, based on promising results of a number of studies.^9,10^ For the success of control programmes, community support is critical. A scabies control program should address the social dimensions such as stigma.^4^ Therefore, assessment of community perceptions about disease causation, knowledge gaps, attitudes towards prevention, treatment access and practices and health-seeking behaviours are crucial to develop effective control programmes.

## Methods

We conducted a qualitative study to explore knowledge and behaviours related to scabies in the Bijagos Archipelago, Guinea-Bissau. The Bijagós archipelago is a group of more than 80 islands located off the coast of Guinea-Bissau, in West Africa. The study was restricted to the largest island, Bubaque. In these remote islands, small dispersed rural communities are organised in small villages and subsist mainly on agriculture and fishing activities.^11^ The predominant ethnic group, called Bijagós, is characterised by a traditional social system with ancient beliefs and traditions. The Bijagós dialect, Guinea-Bissau Kriol (lingua franca) and Portuguese (official language) are the main languages of the region.^11^

### Study design

We conducted semi-structured interviews with community members, community health workers (CHW), healthcare workers and traditional healers. We supplemented this with an oral questionnaire administered to additional community members.

### Interviews’ topic guide and questionnaire development

The topic guide included four main domains covering knowledge and perceptions about the cause and presentation of scabies, attitudes towards prevention, treatment-seeking behaviours and scabies treatment practices, and knowledge and perceptions about the consequences of scabies in health and life quality. The development of the questionnaire was based on similar studies about NTDs conducted in West Africa^12,13,14,11^. We utilised a mix of both closed and open-questions covering each of the main domains. We piloted the questionnaire with five people in the community to evaluate the comprehension of questions and linguistic barriers. Information from the first four interviews was used to refine the final questionnaire.

### Participants recruitment

As this was a qualitative study no formal sample size calculation was conducted. Sufficient individual semi-structured interviews and questionnaires were undertaken until data saturation was achieved. This occurred when data became redundant in providing new information and potential themes.^15^ Individuals aged more than 18 years were recruited. To obtain four different key informant groups - community members, CHW, healthcare workers and traditional healers - snowball sampling was used for interviewees selection. This sampling strategy is useful when participants are difficult to recruit and consist of asking each interviewee to suggest or invite further participants.

For the orally-administered questionnaire, one in five households were randomly sampled across both the villages and neighbourhoods of the semi-urban area of Bubaque. Utilising methodologies of previous population-based surveys conducted in the islands^16,17^, a household list was generated and used to identify households within each village or neighbourhood. A household was defined as “a group of individuals who are fed from the same cooking pot”.^17^ In each house, the oldest adult member present during the visit answered the questions. One day before the questionnaire administration, visits were conducted to provide information about the study and to obtain verbal consent from village leaders.

## Data collection and analysis

Data were collected in June and July 2018. Prior to both the interviews and questionnaire clinical images of typical scabies skin lesions were shown to participants. Semi-structured interviews were conducted in Portuguese or Kriol by the first author and a trained field assistant. Interviews were audio-recorded using a digital recording device. After transcription, thematic analysis was performed. This process consisted of several steps as follows^18^: familiarisation and immersion in the content; descriptive code generation; code categorisation and creation of potential themes, theme definition and selection of extract examples. Responses to the questionnaire were collected on to a mobile device using Open Data Kit. The qualitative component of the questionnaire was also included in the content thematic analysis. Descriptive statistics methods were used for quantitative analysis using STATA version 15 (StataCorp, Texas, USA).

### Ethical statement

This study was conducted in accordance with the declaration of Helsinki. Ethical approval was obtained from the Comitê Nacional de Ética e Saúde (Guinea Bissau), the LSHTM Ethics Committee (UK) and The Gambia Government/MRC Joint Ethics Committee (The Gambia). Verbal consent was obtained from community leaders. Written informed consent was obtained from all study participants. A signature or thumbprint is considered an appropriate record of consent in this setting by the above ethical bodies.

## Results

### Participants background characteristics

In total 26 interviews were performed with 10 community members, 4 CHWs, 6 healthcare workers (doctors, nurses and pharmacists) and 6 traditional healers. The orally-administered questionnaire was completed by 75 individuals.

The overall mean age of participants was 37.5 years. The main religions were Catholic (38.9%) and Animism (34.7%), and most of the questionnaire participants were of Bijagós ethnicity (86.7%). A minority of participants were illiterate (26.7% in the questionnaire and 19.2% in the interviews). Participants’ demographic characteristics are summarised in table 1.

**Table 1.**
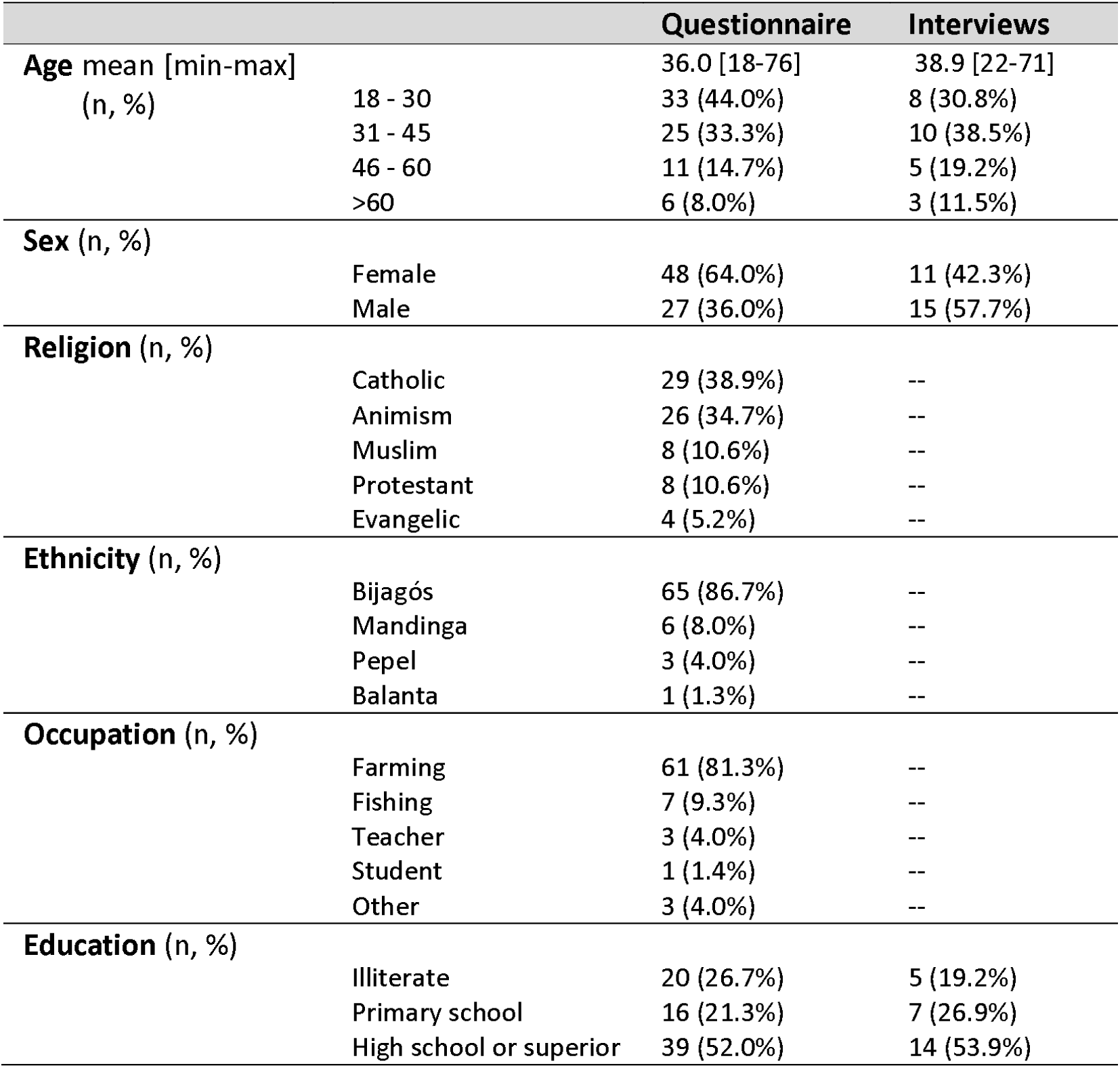
Demographic characteristics.

### Perceptions: Disease awareness

Locally, scabies is informally referred to as coça-coça meaning scratch-scratch or coceira meaning itch. Most participants in the semi-structured interviews reported that scabies is not a common disease compared to other skin diseases, like superficial fungal infections. However, healthcare workers reported that scabies has a significant prevalence, particularly among children under 5 years old. Some healthcare workers reported that scabies could be neglected by the population since it is not considered a severe disease.

Amongst respondents to the questionnaire 96% (n=72/75) reported having heard of scabies infestation but only 42.7% (n=32.75) recognised lesions of scabies when photos with typical skin lesions were shown. Sixty-eight (90.7%) of the questionnaire respondents recognised scabies as an important health problem. A summary of the questionnaire answers about disease perceptions is shown in table 2.

**Table 2.** Main questionnaire answers on perceptions about scabies.

| PERCEPTIONS |  |  |
| --- | --- | --- |
| **Do you recognise the skin disease in the photograph?** | Yes | 32 (42.7%) |
|  | No | 43 (57.3%) |
| **Have you heard about scabies?** | Yes | 72 (96.0%) |
|  | No | 3 (4.0%) |
| **From whom have you heard about scabies?** | Family/friends | 42 (56.0%) |
|  | Healthcare worker | 20 (26.7%) |
|  | School | 13 (17.3%) |
|  | Radio | 5 (6.7%) |
| **How is scabies transmitted?** | I don't know | 29 (38.7%) |
|  | Because of lack of hygiene | 27 (36%) |
|  | Through clothes and bedding | 15 (20%) |
|  | Skin contact with someone infected | 11 (14.7%) |
|  | Through dirtiness | 11 (14.7%) |
|  | Playing on the ground | 6 (8.0%) |
|  | Through mosquito bite | 3 (4.0%) |
|  | Through a mite | 2 (2.7%) |
| **What are the main symptoms of scabies?** | Intense itch | 55 (73.3%) |
|  | Fever | 30 (40%) |
|  | Skin lesions | 26 (34.7%) |
|  | I don't know | 14 (18.7%) |
|  | Pain | 3 (4.0%) |
|  | Malaise | 3 (4.0%) |
|  | Night itch | 2 (2.7%) |
|  | Sleep disturbances | 1 (1.3%) |

### Perceptions: Causation and transmission

Few participants reported that scabies is caused by a mite. The most commonly attributed causes for scabies were dirtiness, lack of hygiene and dirty water. A considerable proportion of participants recognised scabies can be transmitted person-to-person (Table 2).

> “It comes from dirtiness (…), it can pass from one person to another.”.
>
> — (Community member, Int-15)

> “If someone at home has scabies and doesn’t get treatment, he can transmit it to others.”
>
> — (Community member, Int-5)

Some people also believed transmission occurred through contaminated personal items (fomites), namely shared clothes and bedding.

> “You can get scabies if you wear a shirt of someone who is infected.”
>
> — (Community member, Int-7)

Very few participants reported disease causation due to contact with contaminated animals or mentioned a supernatural cause for scabies. Intense itch, fever and skin lesions were the most common reported disease manifestations (Table 2).

### Perceptions: Social impact, stigma and health consequences

Most participants reported intense itching as a reason for inadequate rest with implications on work productivity and school absenteeism.

> “[Children] can’t go to school because of the itching; the teacher teaches (…) but children are constantly scratching the body and don’t learn.”
>
> — (Traditional healer, Int-19)

Moreover, scabies was reported to contribute to social isolation and inhibition, especially among children, who change behaviours and social interactions.

> “…[children] stay alone at home.”
>
> — (Community member, Int-4)

> “the child does not like being with friends, does not feel well.”
>
> — (Community member, Int-5)

There was a general perception that people affected by scabies suffer from stigma. Feelings of rejection, embarrassment and shame were frequently reported and coping strategies to deal with it were mentioned.

> “Adults use strategies to hide the disease. They use long-sleeve shirts and trousers.”
>
> — (Healthcare worker, Int-2)

> “Actually people feel embarrassment (…) people seek for the traditional healer because he works alone. In the hospital people are exposed (…).”
>
> — (CHW, Int-8)

The relation between poverty and scabies was clearly identified with economic problems as both a cause and a consequence of scabies.

> “Economic problems can worsen this kind of situations (…) The treatment-related costs affect the family income. The trip to the hospital implies a loss of daily earning.”
>
> — (Healthcare worker, Int-1)

Only one healthcare worker reported that complicated scabies sometimes needs to be treated with antibiotics and no healthcare workers reported knowledge of post-streptococcal complications or their association with scabies.

### Attitudes: Prevention and health education

Personal hygiene and avoidance of contact with dirty personal items was the principal method for scabies prevention that emerged from both the questionnaire responses and the semi-structured interviews. Questionnaire responses towards attitudes for scabies prevention are listed on table 3.

**Table 3.** Questionnaire responses about attitudes on scabies.

| ATTITUDES |  |  |
| --- | --- | --- |
| **How can you prevent scabies?** | Keep personal hygiene | 69 (92%) |
|  | Keep clothes cleaned | 37 (49.3%) |
|  | Infected people should be treated | 9 (12.0%) |
|  | Keep home hygiene | 9 (12.0%) |
|  | Avoid contact with dirty water | 7 (9.3%) |
|  | Eat safe food | 5 (6.7%) |
|  | Avoid skin contact with infected people | 3 (4.0%) |
|  | Avoid playing on the ground | 3 (4.0%) |
| **If someone in your household is infected with scabies, what do you do to prevent the spread of the disease?** | Infected people should be treated | 61 (81.3%) |
|  | Avoid skin contact with infected people | 11 (14.7%) |
|  | Wash clothes | 9 (12.0%) |
|  | Treat everyone in the household | 2 (2.7%) |
|  | Topical application of a home remedy | 2 (2.7%) |
|  | Don't share the bed with infected people | 2 (2.7%) |

> “To prevent scabies or other skin diseases people need to have hygiene; should always be clean.”
>
> — (Community member, Int-5)

In the same vein, hand hygiene, house cleanliness, avoiding playing on the dirt, clean clothing and soap use were suggested as measures to prevent scabies.

Most participants perceived scabies as a communicable disease by skin-to-skin contact, but few clearly suggested prevention measures such as avoiding personal contact with infected persons. Nonetheless, some participants identified treating those infected as soon as possible as a prevention measure. Moreover, individual treatment was the main questionnaire answer as a measure to prevent scabies transmission within the household (81.3%, n=61).

> “If someone gets scabies, should be treated. Otherwise, their relatives will get scabies if they stay together.”
>
> — (Community member, Int-4)

CHW and healthcare professionals have strong opinions about the role of health education. CHW believe that scabies prevalence is decreasing due to increased levels of hygiene.

> “I think our principal role is on health education and awareness to seek treatment earlier.”
>
> — (Healthcare worker, Int-17)

### Practices: Care-seeking behaviours

Whilst most community members (93.3%, n=70) undertaking the questionnaire reported that they would seek care through the local health centre (table 4), community members participating in interviews reported other treatment seeking patterns including that they would seek care either from a traditional healer or from the health centre. Participants living in villages distant from the health centre were more likely to initially seek treatment from the nearest traditional healer. Healthcare workers believed that first consulting the traditional healer and later the health centre is a common practice.

**Table 4.** Main questionnaire findings on practices about scabies.

| PRACTICES |  |  |
| --- | --- | --- |
| **Where do you seek treatment if someone in your household is infected?** | Local health centre | 70 (93.3%) |
|  | Traditional healer | 7 (9.3%) |
|  | Local pharmacy | 2 (2.7%) |
|  | Homemade remedy | 2 (2.7%) |
| **When do you seek treatment if someone in your household is infected?** | Immediately | 72 (96%) |
|  | Within less than a week | 2 (2.7%) |
|  | Within more than a month | 1 (1.3%) |
| **Which factors influence people's treatment choices?** | Cost of treatment | 66 (88%) |
|  | Cost of transport | 11 (14.7%) |
|  | Trust in traditional medicine | 8 (10.7%) |
|  | Trust in western medicine | 8 (10.7%) |
|  | Distance | 7 (9.3%) |
|  | Believe that scabies goes away by itself | 3 (4.0%) |

> “Sometimes people start treatment with traditional medicines in the village (…) But if you see it doesn’t solve, you decide to go to the hospital.”
>
> — (Community member, Int-22)

Cost was the primary determinant of care seeking (88%, n=66) for most participants. Distance to the health centre and indirect costs related to transportation were other important factors whilst personal preferences for the type of treatment (for example health centre or traditional healer) were also mentioned. Several respondents trusted more in modern medicine and expressed distrust regarding traditional practices.

> “Here [in the village], people think that the traditional healer can help. But no, he can do nothing. Only at the hospital.”
>
> — (CHW, Int-3)

Late presentation, in particular by adults, was reported by both healthcare workers and traditional healers. Community members identified lack of money, but also embarrassment and stigma about the disease as reasons for delayed diagnosis.

> “They seek for treatment at an advanced stage (…). If scabies killed, people would seek care earlier.”
>
> — (Healthcare worker, Int-1)

> “people believe that it can cure by itself (…) when it complicates, they go to the hospital.”
>
> — (Healthcare worker, Int-2)

### Practices: treatment practices

Benzyl benzoate lotion is the only available treatment for scabies in the islands. The most commonly reported types of traditional treatments were leaves or root infusions that could be used for body washing or to drink. Most traditional healers used medicinal plants whilst a minority of them treated patients through rituals and ceremonies. Homemade remedies made from ashes, palm oil and palm fruit were also reported among older villagers.

Treatment of household contacts is not commonly practiced by traditional medicine or allopathic medicine practitioners.

> “We only treat what we see. But normally we should treat everyone in the household (…) We can’t [treat]. We only treat those who came to the hospital.”
>
> — (Healthcare worker, Int-1)

Apart from topical treatment, healthcare workers also recommended that clothing and bedding be washed and exposed to sunlight.

> “People should change clothes on a daily basis (…) Clothes should be washed and dried at the sun to kill the pathogenic germ.”
>
> — (Healthcare worker, Int-2)

## Discussion

This study has explored the Bijagós community perceptions about aetiology, mode of transmission, disease burden and social-economic impacts of scabies; attitudes regarding prevention; and practices concerning treatment-seeking behaviours and scabies treatment.

Community awareness of scabies is considerable. Most participants had heard about scabies, the local terms for scabies were widely recognised, and more than 40% of questionnaire respondents identified the disease from clinical images. The study was undertaken five months after a skin infections prevalence survey in some of the same communities and this could have contributed to the increased recognition of skin diseases.

Despite widespread awareness, a wide range of misconceptions regarding disease causation and modes of transmission was found. Lack of personal and environmental hygiene were commonly believed to be the causes of scabies. This is concordant with the results of qualitative research about other skin NTDs performed in West Africa.^12, 13^ Personal hygiene and environmental cleanliness were believed by respondents to be crucial interventions to prevent scabies. Unlike other knowledge, attitudes and practices (KAP) studies concerning other skin NTDs^12,19^ such as Buruli Ulcer and Yaws, only a minority of participants reported beliefs in supernatural causes for scabies. This might be explained by the different cultural, religious and ethnic background between the studies’ populations. In previous qualitative studies performed in the Bijagós Islands, perceptions about aetiology and transmission of trachoma were more related to lack of hygiene and supernatural causes were not mentioned^11^. There was an accurate perception of transmission by skin-to-skin contact with an infected person and of the need to treat the infected person to prevent scabies within a household. Apart from the itching and skin lesions, community members frequently mentioned fever as a common symptom. Fever could occur when scabies is complicated with a secondary bacterial infection. In the community, the concept of fever seems to be broader than a physiological increased of body temperature. The term “fever” was frequently used as a synonym for feeling unwell.

Whilst treatment of contacts is crucial for scabies control^20^, treatment of contacts was rarely reported by healthcare workers due to logistical challenges in reaching family members and cost implications. Community members do not undertake this additional control measure because treatment is unaffordable. Knowledge about complications of scabies was limited among healthcare workers. Whilst a third of healthcare workers reported the need for concurrent antibiotic treatment of secondary skin bacterial infections, few recognised the risk of more significant complications. Collectively these findings highlight the need for education and training of healthcare workers about the treatment of close contacts and to inform healthcare professionals about the long-term complications of scabies.

Most study participants perceived that scabies affects all age groups equally and there is an overall perception that scabies prevalence is decreasing over time. In recent years, health education campaigns focused on hygiene and sanitation improvement have been implemented in Bubaque^11^. Concordant with the belief that personal and environmental hygiene play a key role in scabies transmission this work was broadly identified as one contributing factor to scabies’ decreased prevalence. Recently, the islands’ communities have received mass drug administration with ivermectin and albendazole for lymphatic filariasis control which may provide a more plausible explanation for the reduced scabies prevalence^21,22^.

Scabies was perceived to have a negative impact on quality of life, affecting personal health, fitness to work and school attendance, with children particularly negatively affected. Our findings are consistent with a previous study assessing the association of ectoparasitic skin disease with quality of life of Ethiopian schoolchildren.^23^ A study in Brazil showed that scabies is associated with feelings of shame, restrictions on leisure activities, behavioural changes, social exclusion and stigmatisation.^24^ Similar themes were mentioned in the current study. Scabies is a stigmatised condition in the Bijagós community, related to fear of transmission. In this study, discriminatory attitudes, feelings of rejection, social isolation and treatment-seeking behaviours were associated with stigma. Late presentation and its health consequences may be influenced by the stigmatising nature of the condition.

There was discordance between the interviews and the questionnaire responses about reported treatment seeking behaviours. The local health centre was reported to be the first choice by most questionnaire respondents, but the more in-depth interviews suggested that community members use homemade remedies or seek for traditional medicine as a first choice. This discordance may be partly explained by social desirability bias on the part of questionnaire respondents. Expectable analogies between the author that administered the questionnaire and western medicine were unavoidable. It may have contributed to biased answers in favour of modern medicine instead of traditional practices. Other factors related to data-collection may also have played a role. Semi-structured interviews were almost always conducted in private, but during questionnaire administration, the presence of other community members was frequent.

The primary determinants of care seeking were reported to be cost and distance to the healthcare facility. Individuals living at a greater distant from the health centre were more likely to report using traditional healers for scabies treatment. This trend has previously been observed in a KAP study on trachoma in Bubaque.^11^ Whilst patients reported a preference for western-medicine many still reported seeking care from a traditional healer. This could be due to factors outlined above but also the costs with accessing medical care at a health facility. Late presentation was frequently reported by healthcare workers, CHWs and traditional healers during the interviews and appeared to be related to beliefs regarding stigma, the relative importance of scabies and a fear of unaffordable costs of treatment.

Based on our findings, scabies control programmes should focus on health education of the community and training of healthcare professionals to promote early treatment and treatment of contacts. Informed communities are crucial to decrease scabies stigmatisation and late presentation. CHW may have a role in improving health education, access to care and to create bridges between traditional medicine and western medicine for a healthier population.

Some authors advocate for the integration of NTD control programmes, particularly skin NTDs.^1^ In settings similar to the Bijagós Islands, there are opportunities for integration of scabies mapping, MDA and surveillance in other ongoing control programmes, such as lymphatic filariasis. Moreover, there is the need to improve disease management and access to affordable treatment. Public health approaches simultaneously targeting several common skin diseases, such as scabies and impetigo, deserve further research. Our work also highlights the importance of systematically assessing the quality of life impact of less serious but more prevalent skin diseases.

This study has several limitations. Firstly, the island of Bubaque is not completely representative of all the islands in the archipelago. Bubaque is better connected to the mainland, and the main health facility of the archipelago is located there. Future studies should ideally include other islands of the Bijagós archipelago. Additionally, the multi-ethnic population found on the mainland of Guinea-Bissau is not represented in this study, and consequently, different health beliefs and perceptions could be missed. Ideally focus group discussions could also have been included as an additional measure to triangulate the findings of the study.

Despite these limitations, this study provided valuable knowledge about health beliefs and community behaviours towards scabies in the Bijagós communities and represents a major contribution to understanding the impact of scabies in endemic communities. Our findings highlight the need to improve health education, recognition, management and affordable access to treatment to support broader scabies control efforts.

## Supporting information

Stroke Check List

## Funding Statement

This work was supported by an MRC Global Challenges Research Foundation Award [grant number MR/P023843/1]. The funders had no role in study design, data collection and analysis, decision to publish, or preparation of the manuscript.

## Notes

#### Summary of Updates

Correction of author name spelling

